# Liam tackles complex multimodal single-cell data integration challenges

**DOI:** 10.1101/2022.12.21.521399

**Authors:** Pia Rautenstrauch, Uwe Ohler

## Abstract

Multi-omics characterization of single cells holds outstanding potential for profiling gene regulatory states of thousands of cells and their dynamics and relations. How to integrate multimodal data is an open problem, especially when aiming to combine data from multiple sources or conditions containing biological and technical variation. We introduce liam, a flexible model for the simultaneous horizontal and vertical integration of paired single-cell multimodal data. Liam learns a joint low-dimensional representation of two concurrently measured modalities, which proves beneficial when the information content or quality of the modalities differ. Its integration accounts for complex batch effects using a tuneable combination of conditional and adversarial training and can be optimized using replicate information while retaining selected biological variation. We demonstrate liam’s superior performance on multiple multimodal data sets, including Multiome and CITE-seq data. Detailed benchmarking experiments illustrate the complexities and challenges remaining for integration and the meaningful assessment of its success.

## 1 Introduction

Single-cell omics technologies have transformed the way we can study cellular systems, and they have quickly become integral to the current biomedical research landscape. Recent advances allow for concurrently measuring multiple modalities, such as gene expression and chromatin accessibility, per cell (Ma et al, 2020a; Lee et al, 2020; Peng et al, 2020). These paired multimodal assays promise unprecedented insight into cellular diversity and relationships between molecular layers and regulatory processes. However, novel data types also pose new and complex challenges for data integration. We need to combine data from distinct modalities providing complementary information, each with unique dimensionality and statistical properties (vertical integration). It is unclear whether it is advantageous to analyze the different modalities separately or at what level of abstraction it may be beneficial to project data into a joint latent space. At the same time, intricate study designs and meta-analyses require sophisticated non-linear batch effect removal (horizontal integration) (Argelaguet et al, 2021).

Solutions for horizontal integration exist for many unimodal data types (Luecken et al, 2022; Cao et al, 2021; Kopp et al, 2022; Ashuach et al, 2022). However, applying these unimodal solutions to multimodal data is not straight-forward, and they showed varying success on complex horizontal integration tasks in a comprehensive benchmark (Luecken et al, 2022). Likewise, several methods for the vertical integration of multimodal single-cell data are available, but most do not explicitly allow for and evaluate complex batch effect removal (Argelaguet et al, 2020; Singh et al, 2021; Hao et al, 2021; Zuo and Chen, 2021; Minoura et al, 2021; Cheng et al, 2022; Duren et al, 2022; Li et al, 2022b). A notable exception is the model totalVI that performs simultaneous horizontal and vertical integration of CITE-seq data using a conditional variational autoencoder (CVAE) (Gayoso et al, 2021). In addition, recent models for the related problem of mosaic integration combine multimodal and unimodal data sets into a single representation (Ashuach et al, 2021; Gong et al, 2021). Though not explicitly designed and tested for paired multimodal data integration, these methods are, in principle, applicable to the task, but are typically not systematically evaluated. The breadth of methods reflects increasingly complex experimental designs and data types in genomics. However, none address and explore all challenges posed by paired multimodal single-cell data, and often, the modeling choices driving the success of the methods remain elusive.

To effectively tackle these challenges, we developed a conditional variational autoencoder-based model for integrating paired multimodal single-cell data. To our knowledge, liam (**l**everaging **i**nformation **a**cross **m**odalities) is the first approach that allows for a principled tuning of the strength of batch mixing via an adversarial loss term that we recently introduced for the horizontal integration of scATAC-seq data (Kopp et al, 2022). In the following, we apply liam to complex experimental designs with replicate data to confidently assess its integration performance. Its early-stage integration strategy leads to higher robustness towards low-quality modalities than later-stage integration. Through ablation studies, we pinpointed the principal contributors to batch integration. At the same time, we demonstrate its competitive performance on multiple distinct multimodal data sets. Our findings demonstrate the need for methods that account for and exploit complex real-world study designs, explore novel avenues for method evaluation, and expose challenges of current benchmarking and evaluation strategies.

## 2 Results

### 2.1 The model liam

Liam (**l**everaging **i**nformation **a**cross **m**odalities) is a model for the simultaneous horizontal and vertical integration of paired multimodal single-cell data. It builds on prior work employing variational autoencoders for dimensionality reduction and horizontal integration of unimodal single-cell data (Lopez et al, 2018; Svensson et al, 2020; Kopp et al, 2022). The deep generative model learns a joint low-dimensional representation of two single-cell modalities while accounting for batch effects. Liam currently supports any pairwise combination of gene expression, chromatin accessibility, and cell surface protein measurements (demonstrated on Multiome and CITE-seq data). We use the negative binomial loss for raw gene expression and CLR-normalized cell surface protein counts and the recently proposed negative multinomial loss for chromatin accessibility data (Kopp et al, 2022).

To enable the model to use potential correlations across modalities at an early stage, the modalities share multiple layers in the encoder that project the data into a shared latent space. To account for intricate study designs resulting in complex batch effects, we model size factors and use a conditional decoder combined with an adversarial training strategy to remove the influence of nuisance variables on the low-dimensional data representation (Ganin et al, 2016; Kopp et al, 2022). The adversarial training strategy introduces a tuneable scaling parameter *α*, with which we optimize the contribution of the batch loss to encourage the mixing of cells from different batches. We employ a logistic-normal distribution for the latent cell variable, making the latent factor loadings interpretable as probabilities (Svensson et al, 2020; Gayoso et al, 2021). We use layer norm for each layer, motivated by the finding of superior horizontal integration performance of the single modality model scVI when using layer norm instead of batch norm (scVI default) (Supplementary note A.1).

Figure 1 shows a complete overview of liam’s architecture. Apart from the default variant of liam developed for paired multimodal data from two simultaneously measured modalities, we also provide variants of liam for unimodal data, e.g., gene expression data (scRNA-seq). We give further details on liam’s implementation in the Methods section.

**Fig. 1.**
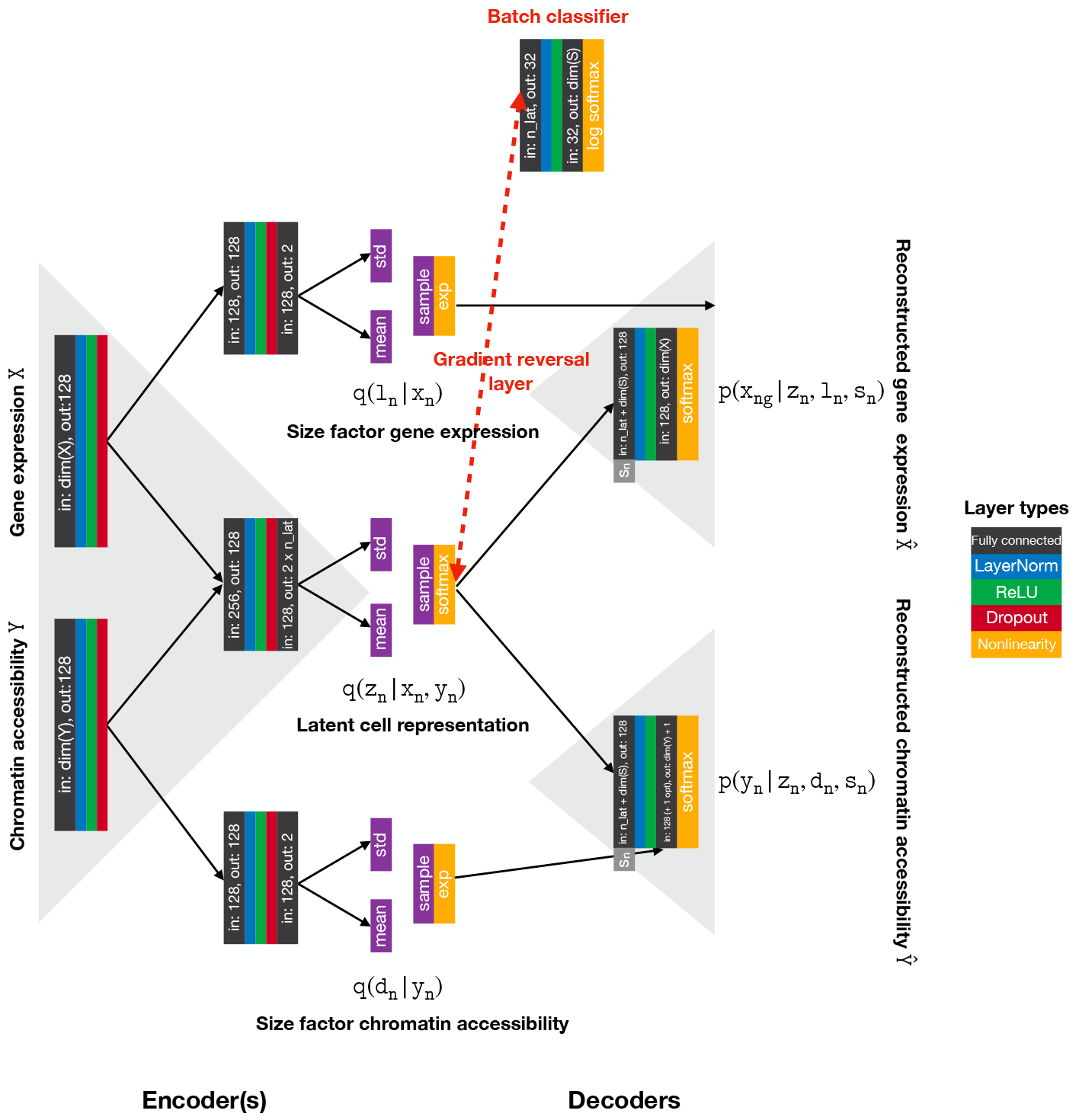
Liam’s architecture. Schematic representation of liam’s architecture for Multiome (paired gene expression and chromatin accessibility data) excluding inferred parameters. The latent cell representation *z*_*n*_ corresponds to the integrated low-dimensional data representation (or embedding, default dimensionality: *k* = 20) that we obtain after training. To remove nuisance variables, we model and infer data type- and batch-specific size factors (*l*_*n*_, *d*_*n*_) and dispersion parameters (not shown), and combine a conditional decoder, where we feed the one-hot encoded batch label of each cell *n* (*s*_*n*_) to the decoder with an adversarial training strategy. The model components required for the adversarial training strategy are labeled in red. The gradient reversal layer is represented by a dashed red line. Representation inspired by Supplementary Figure 13 from (Gayoso et al, 2021).

### 2.2 Exploiting replicates for selected treatment effect retention and enhanced batch mixing

Increasingly complex study designs necessitate flexible integration methods that reduce unwanted noise while retaining biologically meaningful differences. We developed liam such that it can exploit nested batch effect structure and disentangle technical from biological variation. To illustrate this unique feature, we apply liam to a data set for which we have two replicates of a treatment-control experiment (stimulation of T cells; DOGMA-seq use case) amounting to four samples (Mimitou et al, 2021) (cf. Figure 2(f)). This data set comprises measurements of three modalities from the same cell, chromatin accessibility and gene expression (used for model training), and cell surface proteins (used for model validation). Using meta-information collected during sample preparation, we conduct experiments assigning distinct variables as the “batch variable” in the model (i.e., the variable whose effects to remove from the latent representation) (Figure 2). We measure the removal or retention of effects of distinct variables from the collected meta-information with the diversity score iLISI ((Korsunsky et al, 2019), as implemented in scib (Luecken et al, 2022)), for which a higher score indicates better mixing with respect to the chosen variable. When treating the experimental replicate pairs as two distinct batches, liam removes technical variation between replicates while retaining differences between conditions (i.e., treatment and control). By contrast, when treating each sample as a distinct batch, cells from treatment and control samples within and between replicates are mostly mixed, potentially resulting in a loss of biological signal induced by the treatment. Both settings retain known treatment effects, e.g., the emergence of activated T cells marked by the expression of the cell surface protein CD69 after T cell stimulation. However, some more subtle biological differences are only retained by the model penalizing differences between replicates, but not samples, as seen in the split of cell populations by treatment, coinciding with treatment-affected cell surface marker expression not used for training (e.g., the selective depletion of CD3-2 cell surface marker expression in treated samples).

**Fig. 2.**
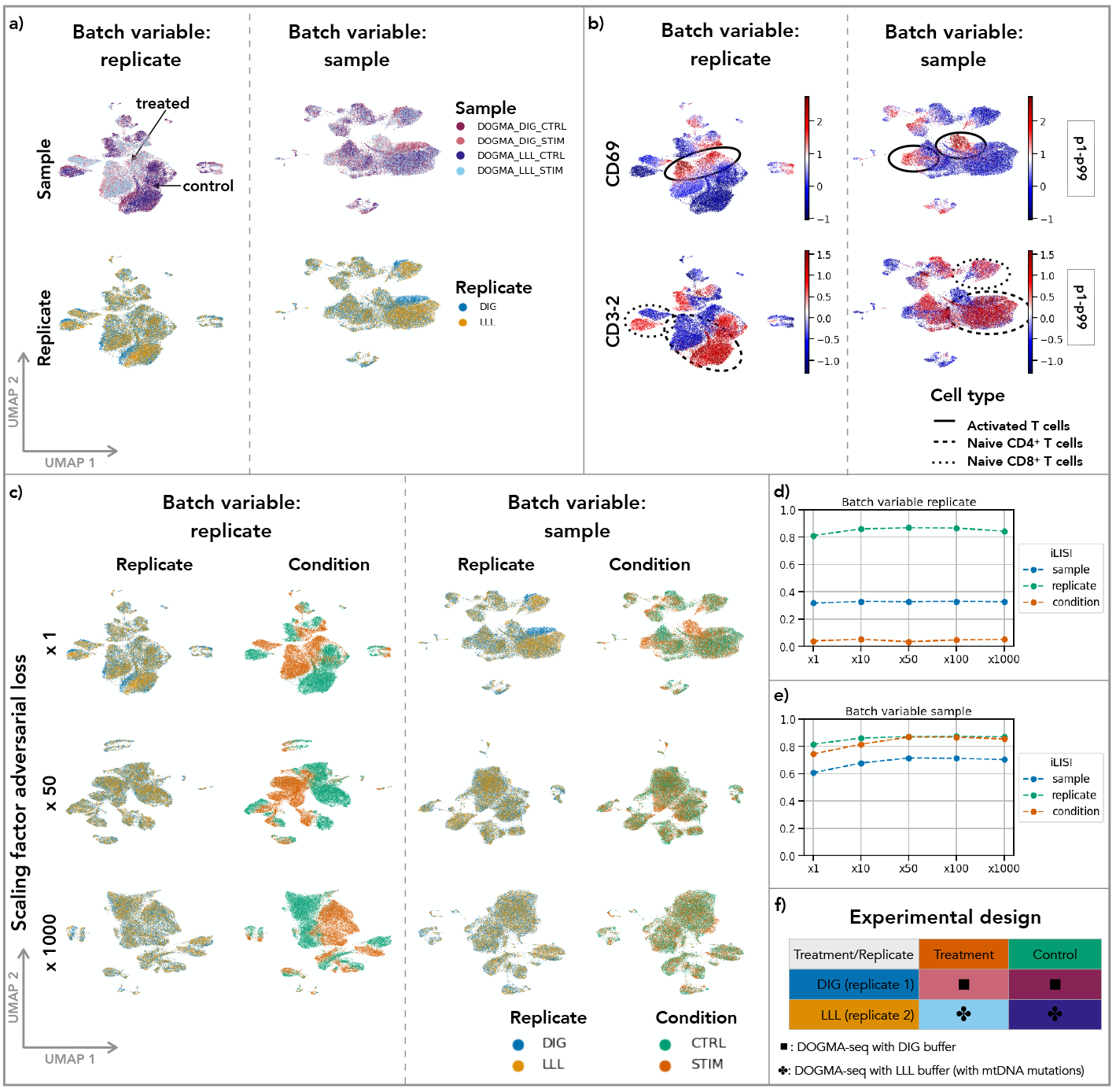
Exploiting replicates, liam preserves selected treatment effects in the data representation and can enhance replicate integration with its adversarial training strategy. Data for all figure panels stems from a treatment/control experiment (T cell stimulation) with replicate (Mimitou et al, 2021). Panel (f) shows the experimental design. Panels (a) and (b) show UMAP representations of the low-dimensional data representations of two models trained with different factors assigned as the batch variable to be removed from the latent representation; one uses “replicate” (left columns) and the other “sample” (right columns). Cells are colored by (a) sample (top) and replicate (bottom), arrows highlight the separation of cells by treatment for the model removing differences between replicates; (b) scaled CLR-normalized cell surface protein counts of the T cell activation marker CD69 (top) and another marker predictive of the treatment, CD3-2 (bottom), different dashed circles highlight cell populations of interest, values outside the p1-p99 percentile range get assigned the min/max value, respectively. (c) UMAP representations of models for which we up-weighted (*α*: x50, x1000) the contribution of the adversarial term in the models loss functions and assigning distinct variables as batch variable; top row corresponds to models in panel (a). (d) and (e) the diversity score iLISI computed for distinct target variables (sample, replicate and condition) for the different variants of liam. The larger the iLISI score, the more mixed samples are with respect to the chosen variable.

These experiments highlight common issues of integrating data sets without replicates: When removing batch effects between samples, we face a tradeoff between preserving biological variation and removing nuisance variables. Assuming that shared variation between multiple replicates profiled under the same condition is more likely to be of biological than of technical origin and that we run a low risk of removing biological signal between replicates beyond sampling variation, we investigate the effect of increasing the contribution of the adversarial loss to the model’s total loss for the previous experiments. In particular, we choose different penalization strengths of batch effects via the scaling parameter *α* using either replicate or sample as the batch variable. Figures 2(c) and (d) demonstrate that we can improve replicate mixing when upscaling the contribution of the adversarial component in liam’s loss function while retaining condition-specific differences ((c) left panel) across a wide range of scaling parameters when choosing replicate as the batch variable. We can optimize this effect via the iLISI replicate metric, which changes with the upscaling of the contribution of the adversarial term. At the same time, the iLISI condition metric stays constant across scaling factors, demonstrating that this approach is robust and does not reduce biologically meaningful variation. When removing all differences between samples (disregarding replicate information, i.e., mirroring the common situation without available replicates), we can also improve replicate mixing (increasing score for iLISI replicate). However, we also remove condition-specific effects when upscaling the contribution of the adversarial term (Figure 2(c) right panel and (e)) (increasing score for iLISI condition). These results underscore the merits of replicates and the power of the batch adversarial strategy implemented in liam concerning the removal of unwanted variation.

### 2.3 Liam excels in a comprehensive benchmark across distinct data types

To further test liam’s capabilities, we use the first-of-its-kind benchmark data set specifically designed to evaluate multimodal single-cell data integration in the NeurIPS 2021 competition “Multimodal Single-Cell Data Integration” (Luecken et al, 2021). To ensure that the data reflects real-world challenges, the organizers generated data from multiple donors across multiple sites, thereby introducing within- and across-site and donor-variation (nested batch effects). This specific experimental design allows for assessing if methods can handle batch effects of distinct sources and scales. We competed with liam for one of the posed NeurIPS competition tasks: “Jointly learning representations of cellular identity” (Task 3). In this competition, liam ranked 4th for Multiome and 2nd for CITE-seq data in the online training category. For this data set, the organizers provide expert-derived cell type annotations as a surrogate for ground truth for cellular state, allowing us to evaluate our modeling choices beyond batch effect removal.

Since the competition metrics were unfortunately either confounded by the nested batch effect structure of the data or had low discriminative power (Lance et al, 2022), we set out for additional evaluations that include unbiased metrics for batch effect removal. In addition to the models that performed best according to the competition’s evaluation criteria for the respective data types and online learning category, we include MultiVI^1^ (Ashuach et al, 2021) for Multiome and totalVI (Gayoso et al, 2021) for CITE-seq data, which are alternative approaches based on VAEs. We trained all models using the sample id (composite of site and donor) as the batch variable, and in contrast to the competition, we did not set any constraints on resource usage.

Figure 3 illustrates that liam is highly effective in removing nested batch effects while retaining biological variation for both Multiome and CITE-seq data (all metrics shown in figures A5 and A6). On Multiome data, liam out-performs MultiVI and LSL AE on the bio-conservation metrics dependent on cell type annotations (nmi and asw label), as well as on trajectory conservation (ti cons batch mean). LSL AE performs best on cell cycle conservation (cc cons), followed by liam and last MultiVI. For evaluating batch effect removal (asw batch and iLISI), we restricted our evaluation to cells from donor 1, for which samples were measured at multiple sites. We assume that batch effects between samples from the same donor should be minimal and mainly reflect sampling and technical differences. Here, liam is best on asw batch d1, and MultiVI on iLISI d1. Both clearly outperform LSL AE, which did not remove the nested batch effects between sites, reflected by the stratification of cells by the site for LSL AE in figure 3(a). For CITE-seq data, liam outperforms totalVI and Guanlab-dengkw on the metric asw label and Guanlab-dengkw on nmi. For the metrics cc cons and ti cons batch mean, Guanlab-dengkw achieves the best results, but the superior performance on those metrics is rendered irrelevant by the model’s failure to integrate data from different sites. As for the Multiome data, liam and totalVI outperform the model performing best in the competition concerning batch integration, with totalVI performing slightly better than liam. In summary, liam successfully removes complex batch effects while preserving biological signal for Multiome and CITE-seq data.

**Fig. 3.**
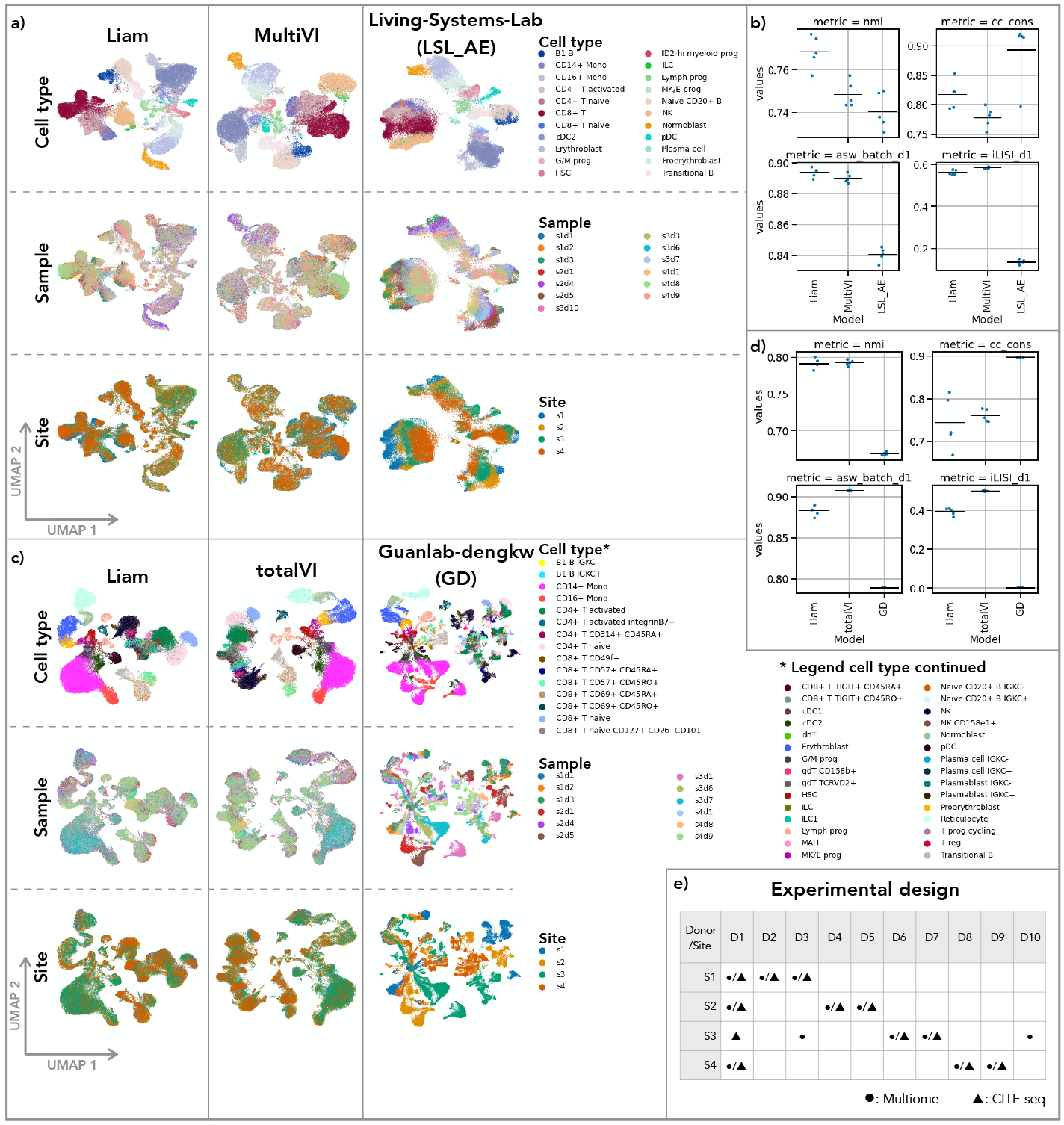
Liam preserves biological variation while removing complex batch effects for multiple data types. Data for all figure panels stems from the NeurIPS 2021 Multi-modal Single-Cell Data Integration competition. Panel (e) shows the experimental design. Panels (a) and (c) show UMAP representations of embeddings obtained with liam and competitors for (a) Multiome and (c) CITE-seq data; Cells are colored by provided cell type annotation (Cell type), sample id (sample) and sequencing site (site). Panels (b) and (d) show selected performance metrics (bio-conservation: nmi, cc cons, batch effect removal: asw batch d1, iLISI d1) with the horizontal line indicating the mean for (b) Multiome and (d) CITE-seq data. All computed metrics, including all competition metrics are shown in figures A5 and A6. For panels (a) and (c) the embedding obtained from a single training run is shown. Panels (b) and (d) show the results of five training runs per model (see Methods).

All of these comparative results need to be considered in the context of the striking observation that a variant of liam, for which we only use RNA as a *single* modality from the Multiome data (cf. Figure A2), performs equally well. Data availability and quality of the individual modalities, the dynamics of the biological system, the granularity of expert annotations, and assumptions behind evaluation metrics may all limit what conclusions can be drawn. While this does not impair our ability to evaluate horizontal integration and to compare our model with alternative approaches, it points to the limit of insights that can be gained from benchmarks, especially concerning minor performance differences. We address this observation in detail in Supplementary note A.3, and we show all models’ performances, including baselines (simpler variants of liam detailed in Methods), in supplementary figures A5 and A6. The issue of annotation is also illustrated by liam readily discovering cell types not present in the competition “ground truth”. When considering the silhouette scores of individual cells with respect to cell type annotations, we observe low silhouette scores in regions in-between cell types. However, we also find a striking example of a low agreement between the reference annotation and obtained clustering for a group of cells annotated as CD8+ T cells. This group of cells expresses the well-characterized MAIT cell markers KLRB1 and SLC4A10 (Park et al, 2019) (Figure A1). Combined with the best concordance with the provided cell type annotations, this suggests that the embedding learned by liam captures the cellular states present in the data sets well.

### 2.4 Examining modeling choices

While the scope of possible evaluations in competitions is limited, the controlled setup of the NeurIPS competition allows us to further dissect modeling choices and gain insights into liam’s strengths. Specifically, we systematically ablate individual model components, with all models using at least batch-specific cell size factors and dispersion parameters (except for VAE, see Methods). The main contribution to the performance is the conditional decoder, with no clear advantage of adding the adversarial component (*α*: x1) or a conditional encoder. The conditional encoder VAE (CEVAE) and adversarial VAE (AVAE; *α*: x1) alone cannot remove batch effects. The performance of batch adversarial VAE (BAVAE, liam default, *α*: x1), conditional VAE (CVAE), and conditional decoder VAE (CDVAE) is indistinguishable (Figures 4(c) and A3).

**Fig. 4.**
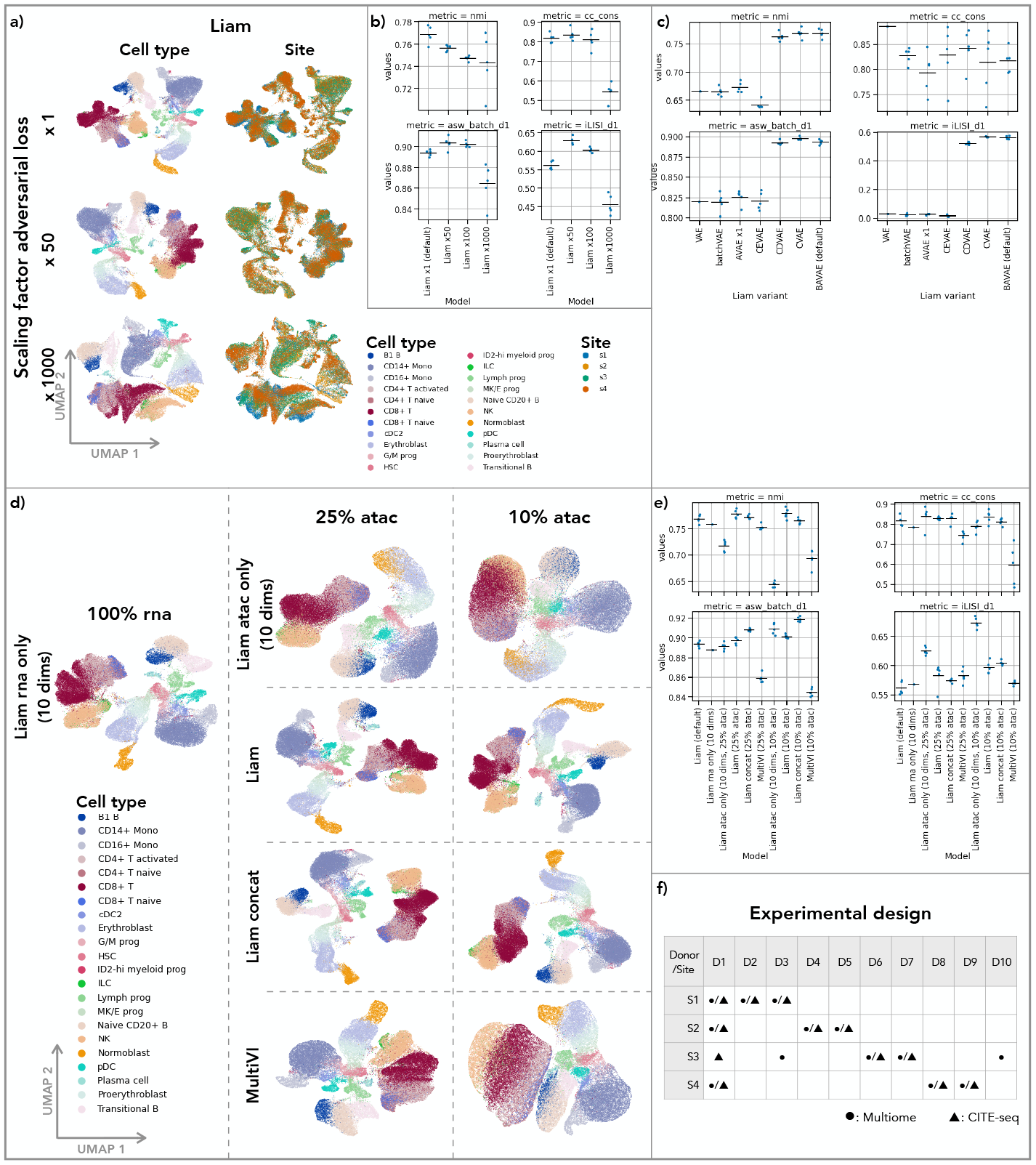
Examining modeling choices exploiting the competition data set. Data for all figure panels stems from the NeurIPS 2021 Multimodal Single-Cell Data Integration competition Multiome data set. Panel (f) shows the experimental design. (a) UMAP representations obtained with liam on competition data set with default setting (*α*: x1) and up-weighting the contribution of the adversarial term by a factor of x50 and x1000; cells are colored by provided cell type annotations and the sequencing site (site). (b) Selected performance metrics (bio-conservation: nmi, cc cons, batch effect removal: asw batch d1, iLISI d1) with the horizontal line indicating the mean. (c) Evaluation of influence of modeling choices on batch effect removal. Performance metrics (as in (b)) for variants of liam with selected model components for batch effect removal ablated. (d) UMAP representations of embeddings obtained with distinct models. Left: a variant of liam trained on rna data only, using the entire rna data. Right: Models using 100% of the rna data and 25% and 10% of tha atac data, respectively. Liam (default, trained on both modalities), Liam concat (concatenation of embeddings from an rna only (10 dims) and atac only model (10 dims)), Liam atac only (10 dims), and MultiVI. (e) Performance metrics (as in (b)) for models in (d) and Liam trained on the full data for reference (Liam (default)). All computed metrics, including all competition metrics are shown in figure A5. For panels (a) and (d) the embedding obtained from a single training run is shown. All other panels show the results of five training runs per model, except for Liam VAE and Liam rna only (10 dims) for which the result of a single training run is shown (see Methods).

When assessing the impact of up-weighting the adversarial contribution in the loss function of liam (BAVAE) (range *α*: x50, x100, x1000), considerable improvements of batch effect removal (asw batch d1 and iLISI d1) are observed, albeit at reduced performance on the cell type label-dependent metrics asw label and nmi. Additionally, while an AVAE with a scaling parameter of 1 fails to integrate the samples horizontally, an AVAE alone can also perform well with a suitable choice for the adversarial scaling parameter (*α*: x50, x100), performing only slightly worse than liam BAVAE with the same scaling parameter choice. Outside a favorable regime for the scaling term (*α*: x1000), liam AVAE appears less stable (Figure A4).

When integrating samples from distinct sources, data quality can vary considerably between samples and, in the case of multimodal data, also between modalities of the same sample. We reasoned that liam may be able to compensate for technical dropouts by exploiting correlation and complementarity between distinct modalities: liam is based on an early-stage joint modeling strategy with a joint encoder architecture, which is in contrast to later-stage integration strategies such as implemented in MultiVI. As the evaluation of liam and the baseline variant liam concat for the high-quality competition Multiome data led to highly similar results (Figure A3(c)), we deliberately decreased the information content of the chromatin accessibility modality by subsampling, to simulate a scenario where we combine a high-quality with a low-quality modality. In particular, we dropped 75% and 90% of the observed features per cell of the binarized ATAC data (this strategy preserves the feature set derived from the full data set). We compared liam to an alternative that concatenates the low-dimensional data representations obtained from two individual modality models (concat) and MultiVI. In this setting, the joint model performs better on bio-conservation than the concat model or MultiVI, with trends reinforcing with diminishing information content (Figures 4 and A5). With diminishing information content, we notice a better mixing of batches for all models (asw batch d1 and iLISI d1), yet again highlighting the trade-off between bio-conservation and batch effect removal. Importantly, liam jointly modeling two modalities performs at least as well as the best performing single modality liam model variant (rna only 10 dims or 20 dims). This finding suggests that liam’s early-stage integration strategy does not impair the model’s performance when jointly modeling a high-and low-quality modality, which we expect to be critical in real-world settings.

## 3 Discussion

Liam is a flexible model for the simultaneous horizontal and vertical integration of single-cell multimodal data. In contrast to other models for multimodal single-cell data, which require the independent horizontal integration of the distinct modalities before vertical integration (Argelaguet et al, 2020; Singh et al, 2021; Hao et al, 2021), liam learns a joint latent representation of two concurrently measured modalities. Currently available multi-omics (benchmark) profiling is strongly biased towards well-studied cell types in human, such as peripheral blood mononuclear cells (PBMCs), where high-quality data has been obtained on different platforms. Recent work on a wider range of cell types and species has shown a broad range of relative data quality and abundance, indicating that laborious case-by-case optimization may be needed to obtain data of similar quality. Liam’s joint representation is demonstrably beneficial when the modalities’ information content and data quality differ and, hence, a much-needed complementary approach. Liam can generally account for complex batch effects, and its adversarial training strategy proves especially useful when replicates are available, enabling the retention of selected treatment effects for treatment/control experiments and the optimal removal of batch effects without running the risk of removing biological variation.

A problem of current multimodal method development is the considerable uncertainty associated with annotations of cellular identity used as a surrogate for ground truth for evaluation. Since they are usually expert-derived from the data itself, they are subject to available knowledge and varying data quality, and they may be biased towards a specific preprocessing strategy and better-studied modalities. For the NeurIPS competition data set, which we chose to illustrate liam’s features, the organizers tried to minimize biasing choices concerning integration approaches and modality. Regardless, the data set is still vulnerable to the problems mentioned, highlighted by the unannotated MAIT cell population in the Multiome data set and the *on par* performance of an RNA-only model with the joint model on the Multiome data (see Supplementary note A.3). To combat this problem, we suggest scrutinizing and updating benchmarks with new insights, considering multiple, independent use cases, and possibly limiting future evaluations to cells that can be reliably annotated across several preprocessing strategies. In general, cell type annotation may not be the most insightful benchmark task for multimodal methods, where the impact of successful integration may be better discernible in downstream tasks such as trajectory inference or network reconstruction (Li et al, 2022a). However, ground truth annotations for such tasks may be even harder to come by or obtain.

Increasingly complex study designs require flexible horizontal data integration strategies. Liam’s adversarial batch effect removal strategy allows for optimizing its strength based on integration metrics across replicates. As this strategy demonstrably retains biological variation, it provides a first-of-its-kind principled solution to the trade-off between technical batch effect removal and biological variance preservation. Future work will focus on testing this framework more thoroughly, on more use cases across species and technologies, and on assessing how to efficiently determine optimal values, e.g., via a simple grid search optimizing batch mixing and monitoring the retention of biological variation. In the absence of replicates or orthogonal data, a user-defined choice of the contribution of the adversarial term is not trivial, as the parameter is likely highly data set-dependent.

We show that liam performs favorably compared to MultiVI in the competition benchmark and when one modality’s data quality is lower. Of note, liam was specifically designed for paired multimodal data, whereas MultiVI was developed with the primary focus of mosaic integration (integration of data sets where some might only contain partially overlapping modalities) and can hence tackle data integration challenges currently not addressed by our model. In this and all other comparisons, we made a deliberate effort to provide fair choices of user-defined parameters, adhering to author-recommended defaults, and testing the robustness to the preprocessing choice of feature preselection for Multiome data (Supplementary note A.2).

In summary, liam provides a flexible, extendable framework for multimodal data integration. Its joint latent space and tuneable batch integration provide demonstrable competitive strengths, making it a method of choice for paired single-cell data.

## 4 Methods

### 4.1 Data sets

#### 4.1.1 NeurIPS competition data set

For the competition use case, we use the phase 2 data of the Multimodal Single-Cell Data Integration NeurIPS competition 2021, accessible through AWS S3. The organizers provide Multiome and CITE-seq data, which comprise gene expression and chromatin accessibility measurements, and gene expression and cell surface protein expression (captured via antibody-derived tags (adt)) measurements, respectively. The data can be retrieved via s3://openproblems-bio/public/phase2-private-data/_joint_embedding/openproblems_bmmc_multiome_phase2/ (Multiome) and s3://openproblems-bio/public/phase2-private-data/joint_embedding/_openproblems_bmmc_cite_phase2/ (CITE-seq). In their study design, the competition organizers purposefully introduced nested batch effects. Their design allows testing generalization capabilities of computational approaches for horizontal data integration by investigating different levels of batch effects removal, e.g., the removal of inter-vs. intra-donor and -site variation. The samples stem from bone marrow mononuclear cells (BMMCs), a complex, disease-relevant, and easily accessible system, and ten distinct donors. The data was generated at four different sites, with samples from one particular donor being measured at four (CITE-seq) and three out of the four (Multiome) sites. For all other donors, a single sample at one site was measured (cf. Figure 3(e)). Each sample is identifiable by a donor site combination (d*s*, with “*” being a wildcard for an identifier for a particular donor and site). The organizers preprocessed and annotated the data from each sample independently. In particular, they preprocessed the distinct modalities separately, deriving independent cell type annotations per modality, which were harmonized afterward into one unified annotation per sample. Of note, the cell type annotations are generally marker gene- or cell surface protein marker-based, including the chromatin accessibility modality, for which the organizers derived gene activity matrices from the chromatin accessibility data before marker gene-based annotation (cf. Supplementary note A.3). A more detailed description of the data set and its preprocessing can be found in appendix A1 of (Luecken et al, 2021).

#### 4.1.2 Treatment/Control data sets from DOGMA-seq

For the treatment/control use case, we use data sets from (Mimitou et al, 2021), which are available on GEO (GSE156478). The data sets are multimodal single-cell data sets of peripheral blood mononuclear cells (PBMCs) that were *in vitro* stimulated with anti-CD3/CD28 and a control (unstimulated). We use data from the DOGMA-seq technology, which measures three modalities at a time, chromatin accessibility, gene expression, and cell surface protein (adt) expression. For DOGMA-seq, a replicate of the experiment is available, as the authors published two DOGMA-seq data sets of the same experiment using two different lysis conditions abbreviated as DIG and LLL. For one of these data sets (LLL), also mitochondrial DNA was profiled, but we do not use this modality here. For the sole purpose of deriving a feature set for the chromatin accessibility data, we also considered samples from the ASAP-seq technology from the same manuscript, which simultaneously profiles chromatin accessibility and cell surface protein levels, as described below. For more information, see (Mimitou et al, 2021).

### 4.2 Data preprocessing

#### 4.2.1 Competition data sets

We used the competition data provided as part of the NeurIPS competition. For ADT counts, we used CLR transformed data (across features) (stored in the field adata.X of the provided AnnData object). For gene expression, we used raw counts, and for ATAC, binarized counts (stored in the field adata.layers[“counts”] in the provided AnnData objects, respectively). The structure of the data set is described in detail in the competition documentation: https://openproblems.bio/neurips_docs/data/dataset/.

#### 4.2.2 Treatment/Control data sets from DOGMA-seq

For the treatment/control use case, we reprocessed the author-provided data. Our reprocessing is loosely based on the preprocessing of the original publication (Mimitou et al, 2021) but was modified to enable the joint analysis of all DOGMA-seq data sets (samples). Each modality underwent separate quality control, and we retain only cells for which all modalities pass it.

##### Chromatin Accessibility

To derive a shared feature set for the chromatin accessibility data from the distinct samples, we jointly analyzed the data from all four DOGMA-seq samples and included chromatin accessibility data from the two ASAP-seq samples from the same study. Starting from the author-provided fragment files, we use an alternative approach to peak calling for feature selection which segments the genome according to cross-cell accessibility profiles called ScregSeg-fi (McGarvey et al, 2022). First, we filter each data set independently using ArchR, only retaining cells with a TSS score exceeding four and a minimum of 1,000 fragments (ArchR version 1.0.0, R version 4.1.2, reference annotation: hg38 from package BSgenome.Hsapiens.UCSC.hg38 version 1.4.1). Next, we remove cells with high counts exceeding Q3 + 1.5x IQR for each data set. Afterward, data from all data sets were combined and used for shared feature calling with ScregSeg-fi. We selected 1,000 bp bins, only autosomes, and only considered regions with at least one count across all cells and binarized the data. We chose the following parameters for ScregSeg-fi: 7 random runs, HMM with 50 states, and 3,000 iterations starting from random initial parameters for each run. As the threshold for informative regions, we chose regions of states that cover at most 1.5% of the genome and that reach a posterior decoding probability of at least 0.9.

##### Gene expression

Starting from author-provided cell-by-feature count matrices, we process each data set separately. First, we remove cells with a total number of unique molecular identifiers (umis) smaller than 1,001. After removing low-count cells, we exclude cells with high umi counts. In particular, those that exceed Q3 + 1.5x IQR. Lastly, we ensure that a minimum of 500 genes was captured per cell and that the percentage of mitochondrial reads is below 30%. As the last step, we exclude mitochondrial genes from the analysis.

##### Cell surface protein expression

Starting from author-provided cell-by-feature count matrices, for each data set, we remove cells that have less than 101 or more than 25,000 counts, that exceed nine control counts, and that have high CD8 and CD4 expression in the same cell. In particular, a cell cannot have more than 30 CD8 counts and 100 CD4 counts at the same time, considering the antibodies for “CD8” and “CD4-1”.

### 4.3 Liam: model and software

#### 4.3.1 Model description

VAEs have been successfully employed for the horizontal integration of unimodal data (Lopez et al, 2018; Svensson et al, 2020; Kopp et al, 2022). Here we modify the prototypical VAE framework, building on recent advances in modeling scRNA-seq data and scATAC-seq data (Lopez et al, 2018; Svensson et al, 2020; Kopp et al, 2022), and introduce liam (**l**everaging **i**nformation **a**cross **m**odalities), a model for paired multimodal single-cell data integration, simultaneously solving the horizontal and vertical integration task.

##### Encoding

Liam’s encoder has two separate input layers for the two modalities of each cell *n*, followed by one hidden layer each. The output of these hidden layers is fed to separate network branches for modeling cell- and modality-specific size factors, with batch-specific priors (*l*_*n*_ for rna and *d*_*n*_ for atac/adt), which are part of our horizontal data integration strategy. Additionally, we concatenate the output of the two modality-specific hidden layers to model the *k*-dimensional (default: 20) latent variable *z*_*n*_, the low-dimensional cell representation, allowing the model to combine information from both modalities.

##### Decoding

We employ two separate decoders, one per modality. These consist of two hidden layers each, which take a sample from the latent variable *z*_*n*_ and the one-hot encoded batch variable *s*_*n*_ (conditional decoder) as input. In this context, “batch” refers to a meta-information variable, such as the condition or sample of a cell. This way, the model can use the batch information for reconstructing the input data without needing to encode batch-related information in the embedding, which is part of our horizontal data integration strategy. We model gene expression and the CLR-transformed adt counts with a negative binomial distribution, using the implementation of Lopez et al (2018) that uses the cell-specific size factor *l*_*n*_. For the chromatin accessibility data, we use the negative multinomial loss, which jointly models a cell’s entire chromatin accessibility profile as introduced in BAVARIA (Kopp et al, 2022), with the modification that we add an extra node to the penultimate fully connected layer, which takes the value of the learned cell-specific atac size factor (*d*_*n*_).

##### Latent factor distributions and inferred parameters

We model cell-specific size factors as log-normally distributed and the latent variable *z*_*n*_ as logistic-normal distributed (Svensson et al, 2020; Gayoso et al, 2021), which has the benefit of the latent factors summing to one, allowing for archetype analysis (Svensson et al, 2020; Gayoso et al, 2021). Additionally, there are several inferred parameters in the model: the batch-specific per gene/adt dispersion of the negative binomial distribution and the batch-specific dispersion of the negative multinomial loss (batch-specific parameters are part of our horizontal data integration strategy).

##### Adversarial training strategy

To further encourage the model to learn latent representations devoid of batch effects and provide a way of tuning batch-mixing, we employ a batch adversarial training strategy. In particular, we introduce an additional neural network, a batch classifier, as a part of our framework that is trained together with the VAE model. The batch classifier has a single fully-connected hidden layer with 32 nodes that takes as input a sample from *z*_*n*_, which is fed through a gradient reversal layer and predicts the batch from which the sample stems. Using a gradient reversal layer allows us to send a negative feedback to the encoder during joint training when the batch classifier gets better at predicting the batch.

##### Further architecture details

All layers are fully-connected layers. We employ dropout layers for the encoder, not for the decoder and batch classifier. We use layer norm and the ReLu activation function for all layers, except for the respective output layers, in which we use other specified nonlinearities (cf. Figure 1). A complete schematic representation of the model’s architecture for Multiome data is shown in figure 1. The figure also details all layer dimensions.

##### Difference between Multiome and CITE-seq architecture

For CITE-seq data, the only difference is that the adt-specific encoder has input dimensions equal to the number of adt features, and that decoder mirrors the rna-specific decoder, with output dimensionality equal to the number of adt features.

### 4.4 Loss function

The model’s loss function comprises regularization terms for the learned latent factors. In particular, we use the Kullback–Leibler divergence for *z*_*n*_, encouraging *z*_*n*_ to follow a logistic-normal distribution (*z*_*n*_ *∼* `Logisticnormal`(0, *I*)), and for the cell-specific size factors of the distinct modalities *l*_*n*_ and *d*_*n*_, encouraging them to follow a log-normal distribution, using the real mean (*l*_*µ*_, *d*_*µ*_) and variance (*l*_*σ*_2, *d*_*σ*_2) of the log of the mean library size per batch (*s*_*n*_) as priors 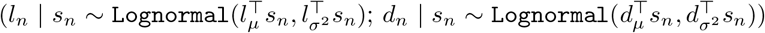. The total regularization loss is:

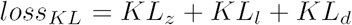

Additionally, the model’s loss function comprises reconstruction loss terms for the distinct modalities. They score the divergence between the input data and the reconstruction with:

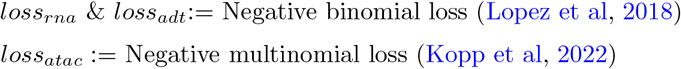

Lastly, the loss function comprises an adversarial term that stems from a batch classifier for which we employ a cross-entropy loss between the predicted class (batch) probability and the real batch (*loss*_*adv*_). In some experiments, we use a user-defined scaling parameter *α*, with which we can up-weight the contribution of the loss of the batch classifier in the total loss (*α* = 1 in liam’s default mode).

We minimize the total loss:

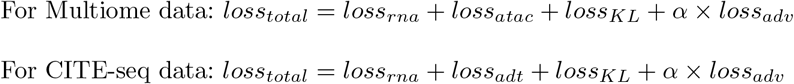

If the batch classifier gets better at predicting the correct batch, this gets fed back to the encoder as negative feedback during the backward pass of model training through a gradient reversal layer (Ganin et al, 2016).

#### 4.3.3 CVAE variant

For the CVAE variant of liam, we remove the batch classifier network. In addition to feeding the one-hot encoded batch variable to the decoder, we feed it to the encoder layers (except for the bottleneck layer) (conditional encoder).

#### 4.3.4 Single-modality model variant

Liam can also be run in a single-modality mode. For this model variant, only the leg of the encoder corresponding to the modality in question is used, and the other is disabled. For the decoder, only the decoder for the respective modality is used.

#### 4.3.5 Model training

For training all variants of liam, we chose a mini-batch size of 128 and split the data into a training set comprising 95% of the data and a validation set comprising 5% of the data. We use Adam for optimizing our model parameters, with a learning rate of 1e-3 and weight decay of 1e-6. We employ early stopping with a patience of ten epochs with respect to the validation loss, using the best model for our analyses. We chose 20 dimensions for the latent space for all models, except for the single-modality models used for the concat baseline, for which we used ten. Models for the competition use case were trained using a Tesla-T4 graphic card with CUDA 11.3. Models for the DOGMA-seq use case were trained using a Tesla-V100-SXM2-32GB graphic card with CUDA 11.3.

#### 4.3.6 Model implementation

Liam is implemented in Python. It employs the scvi-tools library (version 0.14.3) (Gayoso et al, 2022), and we used the scvi-tools-skeleton repository (version 0.4.0) as a starting template for package development. It is available as a readily installable Python package with documentation on GitHub (https://github.com/ohlerlab/liam).

### 4.4 Evaluation

#### 4.4.1 Baselines

##### Horizontal integration

To analyze the contribution of individual modeling choices to the model’s horizontal integration capabilities, we systematically ablate individual model components with all models having batch-specific size factors and dispersion parameters (except for VAE). We compare a:

- VAE: no batch correction at all
- batchVAE: VAE + batch-specific cell size factors and dispersion parameters
- AVAE x1: batch adversarial term only (*α* = 1)
- CEVAE: conditional encoder only
- CDVAE: conditional decoder only
- CVAE: conditional encoder and decoder
- AVAE x1, x50, x100, x1000*; scaled batch adversarial terms
- BAVAE x1 (default), x50, x100, x1000*; scaled batch adversarial terms + conditional decoder

* numbers indicate scaling parameter *α* for the adversarial loss

##### Vertical integration

We compare liam, in which we employ early-stage joint modeling, to simpler baselines. In particular, we compare liam to two single-modality variants of liam with the same dimensionality of the latent space each and a “concat” model. For the concat model, we train two single-modality variants of liam with a latent space dimensionality of half the size of the default model. We concatenate the embeddings obtained from the independently trained single-modality models such that the latent space of the “concat” model space has an equal number of dimensions as liam (default).

- BAVAE rna only (same dimensionality as joint/default; *k* = 20)
- BAVAE atac/adt only (same dimensionality as joint/default; *k* = 20)
- BAVAE concat (concatenation of latent spaces of BAVAE rna + atac/adt only (half dimensionality as joint each; *k* = 10, each)

#### 4.4.2 Other models

##### Models for multimodal data

###### MultiVI

We ran MultiVI (scvi-tools version 0.14.3) with default parameters following a tutorial provided by the authors (https://docs.scvi-tools.org/en/stable/tutorials/notebooks/MultiVI_tutorial.html, as of April 15th, 2022). Of note, this includes a feature preselection step before training the model (also applied in the preprint (Ashuach et al, 2021)). In particular, features present in less than 1% of the cells are removed before training. For comparability with liam, we also run a model without feature preselection. The dimensions of the latent space are determined automatically by the model and are dependent on the number of input features. For the model with feature preselection, the latent space has 16 dimensions, for the model without feature preselection, 18.

###### totalVI

We ran totalVI (scvi-tools version 0.14.3.) with default parameters following a tutorial provided by the authors (https://docs.scvi-tools.org/en/stable/tutorials/notebooks/totalVI.html, as of April 15th, 2022). Of note, this includes total count normalization to 10,000 reads per cell, followed by logarithmization and a feature preselection step before running the model (scanpy version 1.8.2). In particular, for the gene expression modality, the top 4000 highly variable genes were determined with the parameter “flavor: seurat v3”. The latent space dimensionality defaults to 20 dimensions.

###### LSL AE

Best performing model in original competition framework for task 3 Multiome online category according to competition evaluation criteria (Team name: Living-Systems-Lab; method name: LSL AE). We adapted the publicly available code from the submissions to the competition (https://github.com/openproblems-bio/neurips2021_multimodal_topmethods/blob/main/src/joint_embedding/methods/lsl_ae/run/script.py) to be compatible with our analyses. The embedding generated by the model has 64 dimensions.

###### Guanlab-dengkw

Best performing model in original competition framework for task 3 CITE-seq online category according to competition evaluation criteria (Team name: Guanlab-dengkw; method name: Guanlab-dengkw (GD)). We adapted the publicly available code from the submissions to the competition https://github.com/openproblems-bio/neurips2021_multimodal_topmethods/blob/main/src/joint_embedding/methods/Guanlab-dengkw/run/script.py to be compatible with our analyses. The embedding generated by the model has 100 dimensions.

##### Models for unimodal data

###### scVI

We set up two variants of scVI (scvi-tools version 0.14.3). One with default parameters, and one with layer norm instead of the default batch norm for the encoder and decoder. For training, we selected the same user-defined parameter we used for liam.

### 4.5 Stability and subsampling analysis

#### 4.5.1 Treatment/control use case

For the treatment/control use case, we trained one model each with a random seed of 0.

#### 4.5.2 Competition use case

For the competition use case, we compare the performance in the joint integration task of liam to variants of liam and several other models. To account for stochasticity in the training processes, we trained five models each, setting a random state component of the respective frameworks to 0, 994, 236, 71, and 415. All UMAPs show the embedding obtained with a random seed of 0. An exception is the baseline model liam VAE, which was only trained with a random seed of 0.

##### Subsampling analysis

For the subsampling analysis, we fix the random seed of the training process to 0 but introduce stochasticity by using five distinct random subsets of the chromatin accessibility data, dropping 75% and 90% of the binarized ATAC observations each (random seeds for subsampling: 8831, 234, 11, 9631, 94). The UMAPS in the corresponding figures show the embedding obtained with a random training seed 0 using the random ATAC subsample obtained with the seed 8831. This implies that for the concat variant of liam in the subsampling analysis, the 10-dimensional RNA-only data representation is constant (trained with random seed 0), and only the ATAC-only representation has varying input obtained with different random seeds but is trained with a random seed of 0.

#### 4.5.3 Metrics - Competition use case

To evaluate the removal of batch effects and the preservation of biological variation, we use the same six metrics used in the NeurIPS competition. Those metrics are implemented in the scib Python package (comprehensively described in (Luecken et al, 2022)) (scib version 1.0.1). We use the full competition metric name here and denote the shorthand used in the manuscript figures and text in brackets. Four of these metrics score the conservation of biological variance. Two of them depend on cell type annotations provided by the competition organizers, namely NMI cluster/label (nmi) and cell type ASW (asw label). The other two are cell cycle conservation (cc cons) and trajectory conservation (ti cons batch mean). We also use the two metrics scoring batch effect removal from the competition, batch ASW (asw batch sample), using the sample id as the batch variable, and graph connectivity (graph conn). As the metric graph conn was not sufficiently discriminative and the metric asw batch sample using the sample id was confounded by the nested batch effect structure of the data (Lance et al, 2022), we use additional batch removal metrics less prone to confounding. In particular, we use a complementary batch removal metric-iLISI (graph iLISI as implemented in scib v1.0.1 (Luecken et al, 2022)) on the site (iLISI site), and the sample id when subsetting the data to data from d1 (iLISI d1). Additionally, we compute the metric batch ASW on the site (asw batch site) and on sample id when only considering data from donor 1 (asw batch d1). For donor 1, samples were measured at each site (one technical dropout for Multiome data). We compute the batch effect removal metrics only on data from donor 1, as we presume that biological variation between samples from the same donor should be minimal. We reckon that subsetting the data to data from the same donor is a good proxy for scoring technical batch effect removal, not penalizing the retention of potential remaining inter-donor variation.

#### 4.5.4 Metrics - Treatment/Control use case

For quantitatively evaluating horizontal integration success, we computed the iLISI metric (graph iLISI as implemented in scib v1.0.1 (Luecken et al, 2022)) on distinct variables available as meta-information - sample, condition, and replicate (lysis condition). For visualization purposes, we show CLR transformed, scaled ADT counts not used during model training, and raw gene expression values to color cells in UMAP representations.

### 4.6 Code availability

For reproducibility: Analysis scripts https://github.com/ohlerlab/liam_manuscript_reproducibility). Software: Legacy version used for presented analyses: https://github.com/ohlerlab/liam_challenge_reproducibility). Software under development: liam https://github.com/ohlerlab/liam).

## Acknowledgments

We wish to thank Wolfgang Kopp, Leif Ludwig, Remo Monti, Sepideh Saran and Anna Hendrika Cornelia Vlot for critical scientific discussion. We thank Ricardo Wurmus and Mădălin Ionel Patraşcu for technical support. We acknowledge support by the DFG Research Unit FOR 2841 and the DFG International Research Training Group IRTG 2403 and the Chan Zuckerberg Initiative “Single-Cell Biology Data Insights”.

## Appendix A Extended Analyses

### A.1 Layer normalization leads to favorable performance over batch normalization

Early during model development, we tested different normalization strategies in the scVI framework (Lopez et al, 2018) (an RNA-only model) using the DOGMA-seq data set (using only the RNA modality, results not shown). Our analyses suggested that using layer normalization instead of batch normalization (scVI default) leads to better horizontal integration results. This observation led us to use layer normalization in liam. Post hoc, we were able to reproduce this observation with the competition data set, for which we have reference cell type labels available. We compare scVI with its default settings (batch normalization) to a variant of scVI using layer normalization, seeing a clear improvement in the metrics measuring batch effect removal and also for bio-conservation, except for cell cycle conservation (cc cons) (Figure A5).

### A.2 Effect of feature preselection on performance of MultiVI and convergence comparison

For the analyses including MultiVI, we followed the example of the corresponding preprint (Ashuach et al, 2021) and a tutorial (https://docs.scvi-tools.org/en/stable/tutorials/notebooks/MultiVI_tutorial.html) which entail a feature preselection step recommending to use only features present in more than 1% of the cells for modeling. We wanted to rule out that this feature preselection step is a major contributing factor to the observed differences in model performance. Hence, we trained a MultiVI model without feature preselection (abbreviated as fp in figures) but otherwise identical settings. Note that the number of dimensions of the resulting embedding is different, as it is automatically determined by MultiVI and dependent on the number of input features. Figure A5 shows how omitting feature preselection affects the performance metrics. All trends observed with MultiVI with feature preselection are preserved for MultiVI without feature preselection. Liam outperforms both MultiVI variants on bio-conservation and does better on the asw-based batch removal metrics. The MultiVI variants do better on the iLISI-based batch removal metrics.

While MultiVI (Ashuach et al, 2021) models chromatin accessibility data with a Bernoulli distribution, we use a negative multinomial distribution (Kopp et al, 2022). We observe that MultiVI needed substantially more epochs to converge. For liam, the average number of epochs until early stopping was: 54.8 *±* 4.09*σ*, for MultiVI with default settings: 212.4 *±* 27.11*σ*, and for MultiVI no feature preselection: 189.8 *±* 9.47*σ*.

### A.3 Limitations of current benchmarks concerning vertical integration evaluation

As part of our method evaluation, we sought to compare the performance of liam jointly modeling both modalities of the paired NeurIPS competition data sets to variants of liam that use only one of the individual modalities each (baseline). Surprisingly, this obvious baseline crucial for understanding the advantages of a method is rarely computed. Figure A2 shows that the RNA-only model performs on a par with, if not slightly better than the joint model with which we participated in the competition for the Multiome data. Given that the competition was set up to score multimodal data integration, it may appear discouraging that a model trained on a single modality seemed to perform best in the framework. The small difference in performance between the joint and the RNA-only model highlights an important limitation in the expressiveness of the benchmark for evaluating vertical integration: While it is possible that the RNA-only model indeed performs best, another possibility is that the RNA-only model’s superior performance is an artifact of our (RNA) gene expression-centric prior knowledge affecting the definition of cellular states. The cell identity labels used for evaluation (nmi and asw label) were derived independently per data set and modality by the competition organizers and harmonized afterward. Luecken et al (2021) did this in an attempt to capture data set-specific substructure in the final cell type annotations. Regardless, the annotations for the Multiome data set are gene expression-centric, as the chromatin accessibility data was converted to gene activity (GA) scores, thus largely ignoring information from intergenic cell type-specific regulatory elements, and the subsequent annotation is based on known GEX markers. In fact, it has been shown that reducing chromatin accessibility signal to GA matrices before dimensionality reduction results in a substantial information loss (discussed in (Rautenstrauch et al, 2022)). The on par performance of an RNA-only model with the joint model on the Multiome data on cell type label-dependent metrics also suggests that we cannot recover more information on the provided cell type level with a joint than with an RNA-only model for the Multiome data set. For the Multiome data, any existing substructure that a joint model might better recover seems to be less than the overall uncertainty and noise in the labels, leading to all models hitting a maximal performance at around 0.6 for the metric asw label. Pre-trained models seem to have been able to achieve slightly higher scores (Lance et al, 2022), but the incompleteness of the annotations mentioned earlier (e.g., the missed MAIT cell population) let us question the meaningfulness of these results. Additionally, we currently still lack expressive metrics capturing modality-specific information, as the GEX-based cc cons metric (Rautenstrauch et al, 2022), on which the ATAC-only model performed notably worse. This is in line with Ma et al (2020b), who observed a less localized cell cycle signature in an embedding derived from scATAC-seq data compared to an embedding derived from scRNA-seq data when treating modalities from paired data from the same cell independently.

The observed limitations are alleviated slightly for CITE-seq data, where the joint model performed better on capturing the harmonized reference cell type annotations. This is in line with previous observations of some populations being better discernible with ADT than GEX and vice versa, but may be different for other biological systems with less well defined cell surface markers and antibody availability.

We tried to circumvent these limitations by exploring alternative ways to use the benchmark data, simulating challenging (real-world) conditions of differing coverage across modalities to test different modeling choices. Our results suggest that, in practice, joint modeling may be beneficial. In any case, all our model variants do well on horizontal integration and seem to capture the overall biological information contained in the data, especially the RNA-only model for Multiome data, and the joint models. It will be interesting to use the embeddings derived with liam as a starting point to explore one of the main benefits of the paired data, ground truth on relationships between modalities.

---

**Fig. A1.**
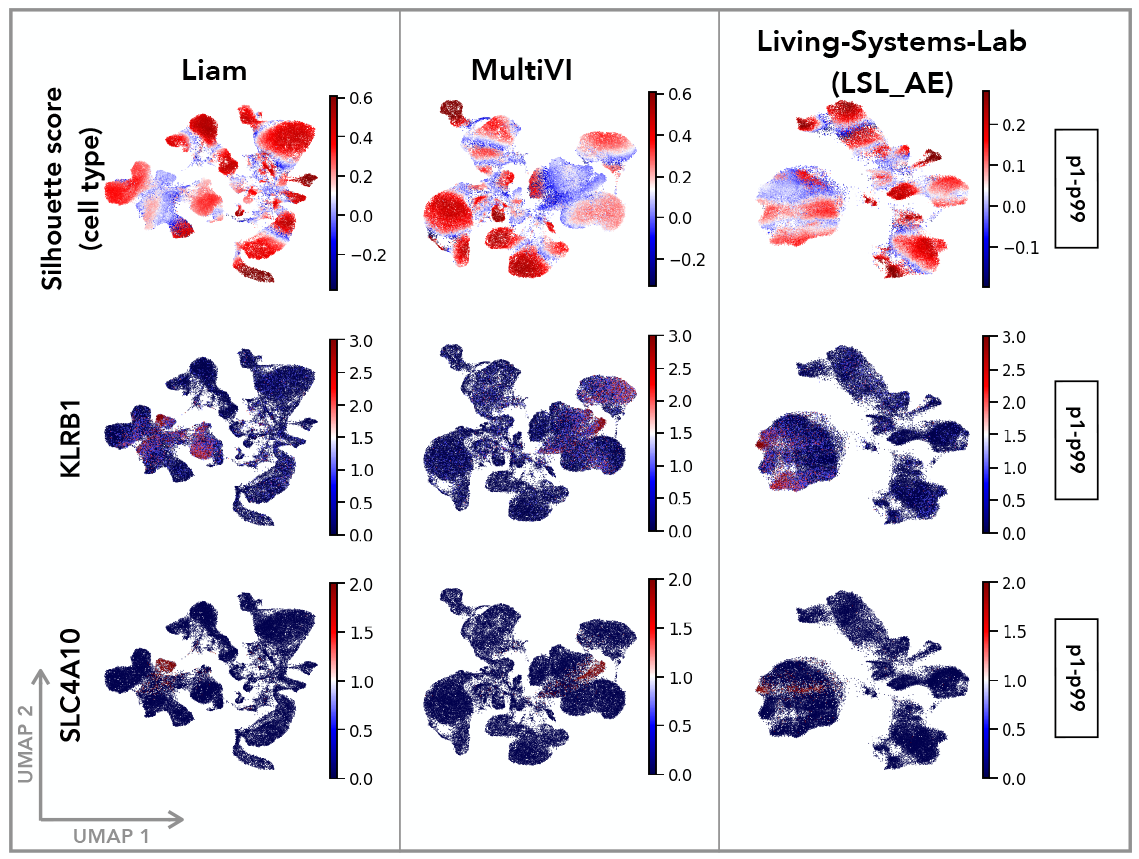
Liam captures cellular subpopulations missed in the NeurIPS competition reference annotation. Data for all figure panels stems from the NeurIPS 2021 Multimodal Single-Cell Data Integration competition Multiome data set. Shown are UMAP representations of the embeddings obtained with liam, MultiVI and LSL AE with cells colored by - first row: per cell silhouette scores with respect to provided cell type annotations; second and third row: raw gene expression values for the MAIT cell markers KLRB1 and SLC4A10 (Park et al, 2019), values outside the p1-p99 percentile range get assigned the min/max value, respectively. The embedding obtained from of a single training run is shown (see Methods).

**Fig. A2.**
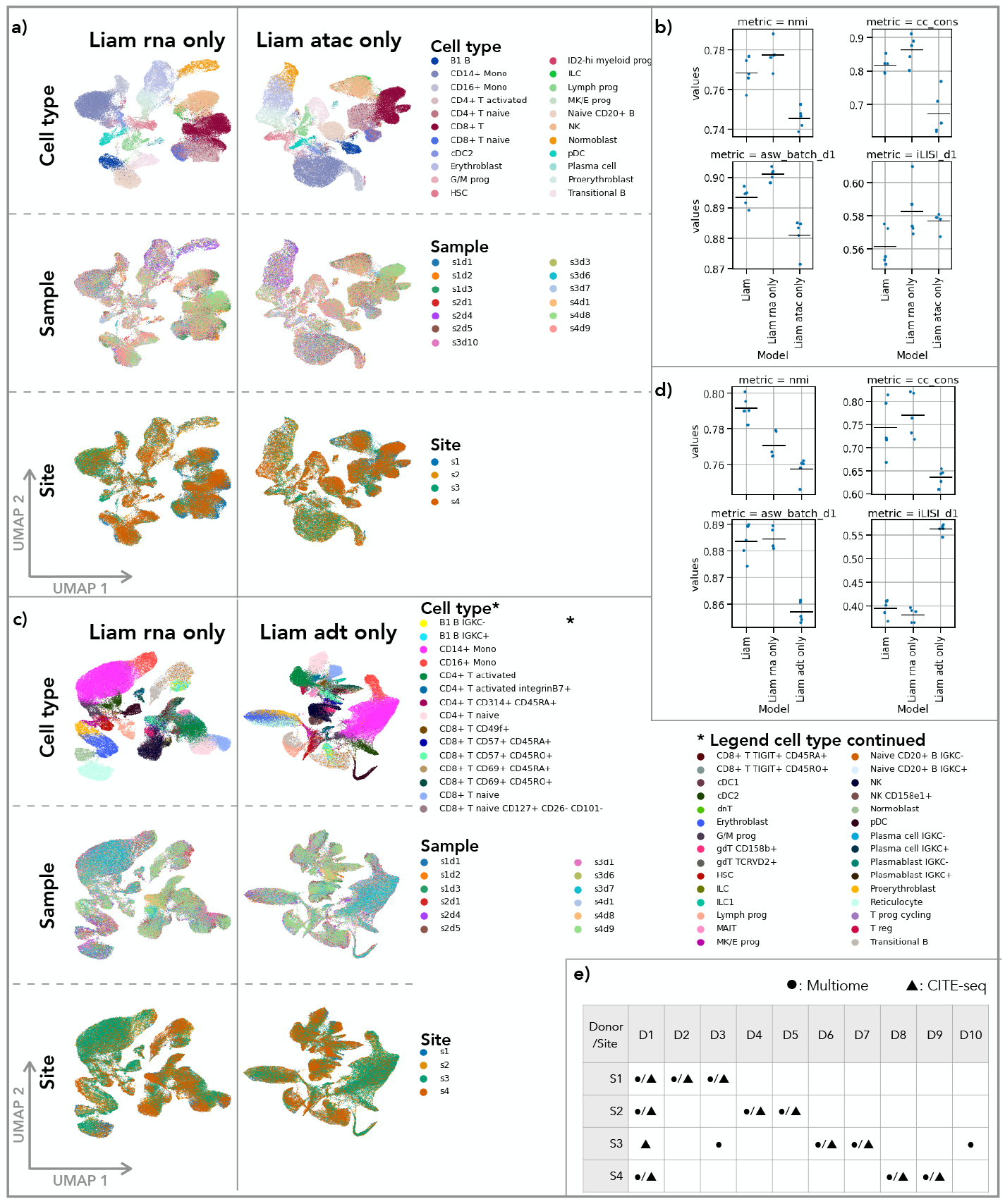
Performance of variants of liam using only single modalities of the multimodal data from the NeurIPS competition framework. Data for all figure panels stems from the NeurIPS 2021 Multimodal Single-Cell Data Integration competition. Panel (e) shows experimental design. Panels (a) and (c) show UMAP representations of embeddings obtained with variants of liam using the individual modalities of (a) Multiome and (c) CITE-seq data, respectively; Cells are colored by provided cell type annotation (Cell type), sample id (sample) and sequencing site (site). Panels (b) and (d) show selected performance metrics (bio-conservation: nmi, cc cons, batch effect removal: asw batch d1, iLISI d1) with the horizontal line indicating the mean for (b) Multiome and (d) CITE-seq data. All computed metrics, including all competition metrics are shown in figures A5 and A6. For panels (a) and (c) the embedding obtained from a single training run is shown. Panels (b) and (d) show the results of five training runs per model (see Methods).

**Fig. A3.**
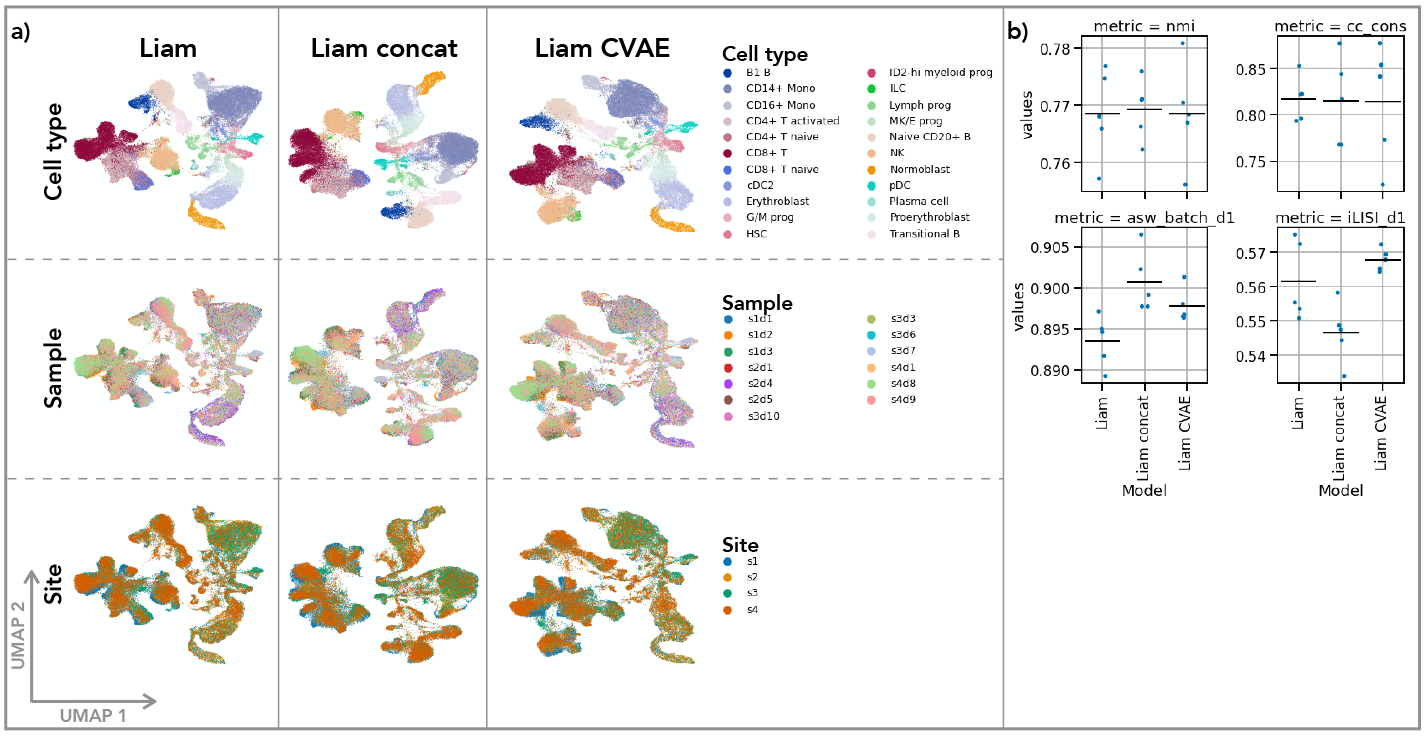
Comparable performance of liam and baseline variants of it on the competition Multiome data. Data for all figure panels stems from the NeurIPS 2021 Multimodal Single-Cell Data Integration competition Multiome data set. Panel (a) shows UMAP representations of embeddings obtained with distinct variants of liam; Cells are colored by provided cell type annotation (Cell type), sample id (sample) and sequencing site (site). Panel (b) shows selected performance metrics (bio-conservation: nmi, cc cons, batch effect removal: asw batch d1, iLISI d1) with the horizontal line indicating the mean of those models. All computed metrics, including all competition metrics are shown in figure A5. For panel (a) the embedding obtained from a single training run is shown. Panel (b) shows the results of five training runs per model (see Methods).

**Fig. A4.**
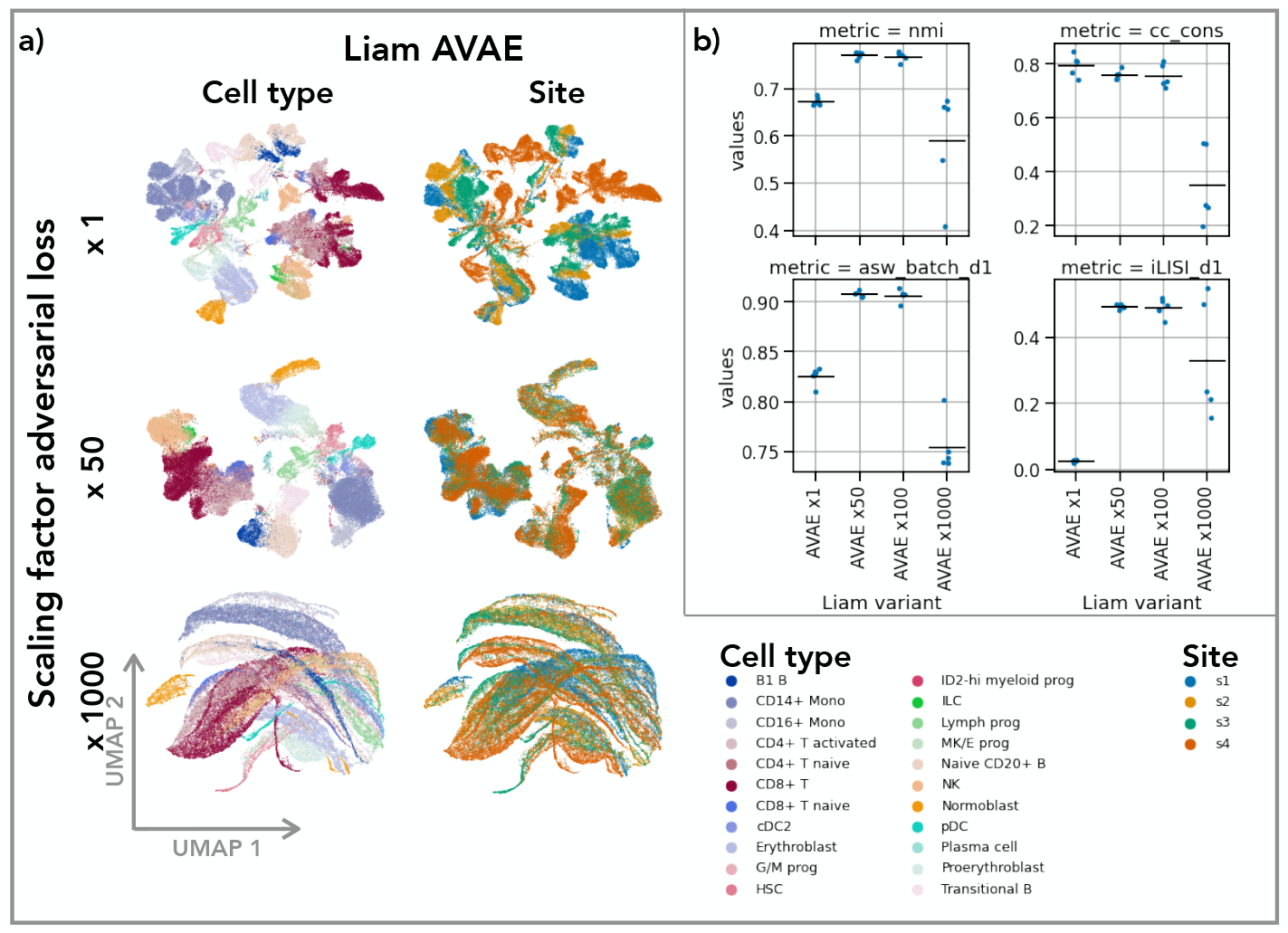
Adversarial training alone (liam AVAE) can perform well on the competition data set for suitable adversarial scaling paramters. Data for all figure panels stems from the NeurIPS 2021 Multimodal Single-Cell Data Integration competition Multiome data set. (a) UMAP representations obtained with liam AVAE with distinct contributions of the adversarial term to the overall loss function (*α*: x1, x50, x1000); cells are colored by provided cell type annotations and the sequencing site (site). (b) Selected performance metrics (bio-conservation: nmi, cc cons, batch effect removal: asw batch d1, iLISI d1) with the horizontal line indicating the mean for liam AVAE models. All computed metrics, including all competition metrics are shown in figure A5. For panel (a) the embedding obtained from a single training run is shown. Panel (b) shows the results of five training runs per model (see Methods).

**Fig. A5.**
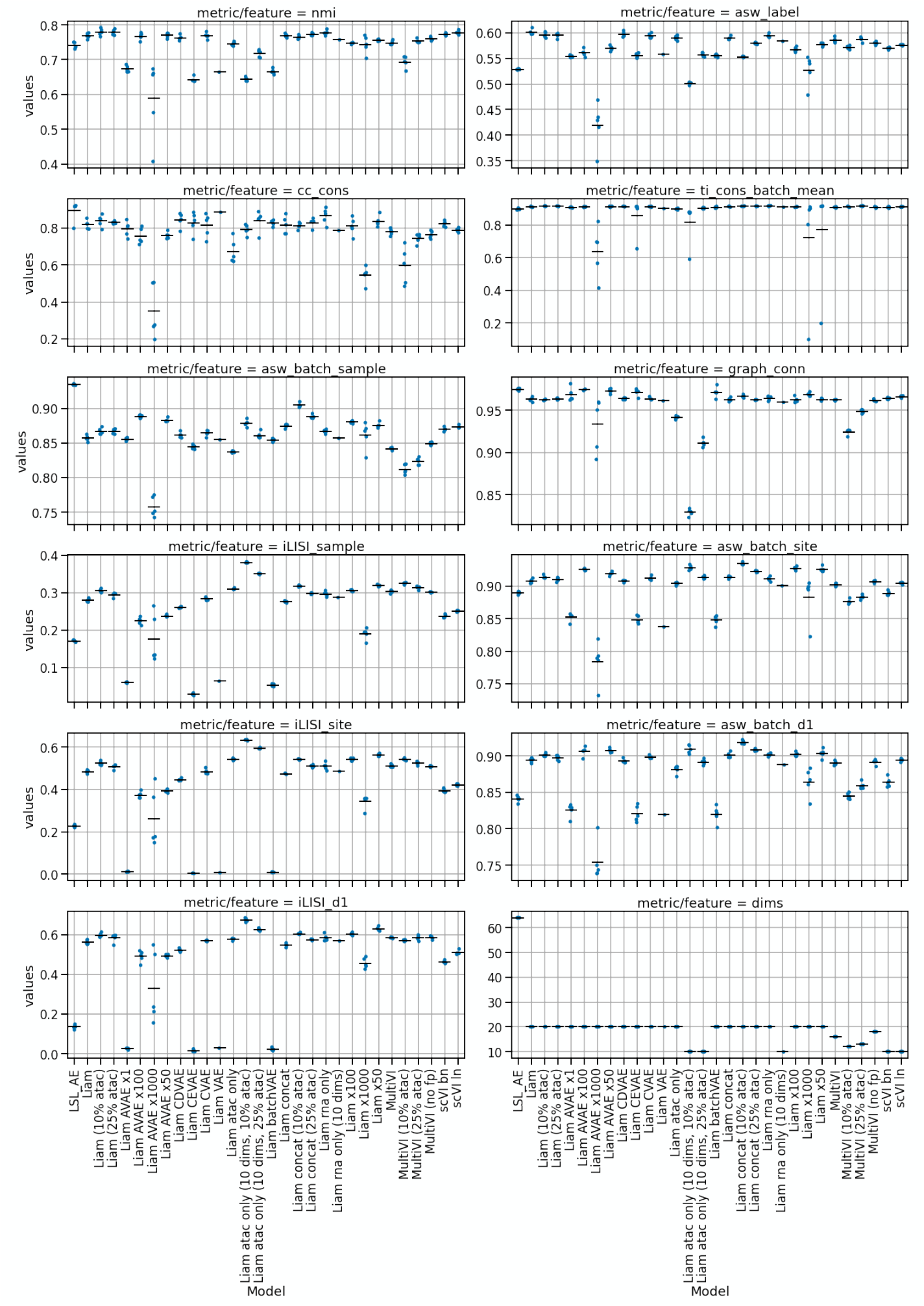
All evaluation metrics for all presented models and the NeurIPS 2021 Multimodal Single-Cell Data Integration competition Multiome data set. Shown are the results of five training runs per model, except for Liam rna only (10 dims) and Liam VAE for which the result of a single run are shown (see Methods). The horizontal line indicates the mean.

**Fig. A6.**
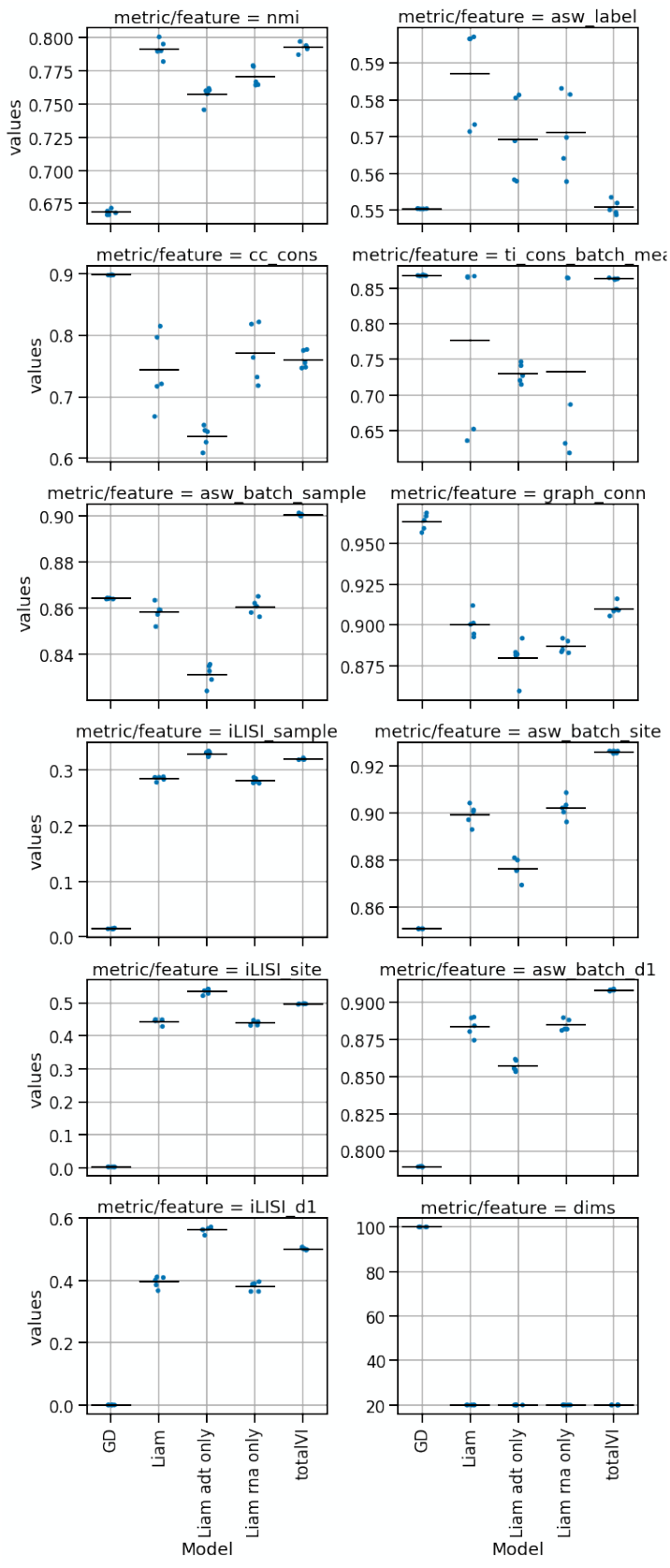
All evaluation metrics for all presented models on the NeurIPS 2021 Multimodal Single-Cell Data Integration competition CITE-seq data set. Shown are the results of five training runs per model; the horizontal line indicates the mean. GD: Guanlab-dengkw. Shown are the results of five training runs per model (see Methods).

## Footnotes

1 The model MultiVI is designed for mosaic integration, of which the paired multimodal data integration studied here is a subtask.

